# Arterial cells support the development of human hematopoietic progenitors in vitro via secretion of IGFBP2

**DOI:** 10.1101/2022.10.04.510611

**Authors:** Paolo Petazzi, Telma Ventura, Francesca Paola Luongo, Alisha May, Helen Alice Taylor, Nicola Romanò, Lesley M. Forrester, Pablo Menéndez, Antonella Fidanza

## Abstract

Hematopoietic stem and progenitor cells develop from the hemogenic endothelium located in various sites during development, including the dorsal aorta from where Hematopoietic Stem Cells (HSCs) emerge. This process has proven especially challenging to recapitulate *in vitro* from pluripotent stem cells and further studies are needed to pinpoint the missing stimuli *in vitro*. Here, we compared iPSC-derived endothelial cells and *in vivo* HSC-primed hemogenic endothelium and identified 9 transcription factors expressed at significantly lower levels in cells generated *in vitro*. Using a novel DOX-inducible CRISPR activation system we induced the expression of those genes during *in vitro* differentiation. To study the phenotypical changes induced by the activation of target genes, we employed single cell RNA sequencing in combination with engineered gRNA that are detectable within the sequencing pipeline. Our data showed a significant expansion of arterial-fated endothelial cells associated with a higher *in vitro* progenitor activity. The expanded arterial cluster was marked by high expression of *IGFBP2* and it was distinct from the hemogenic cluster that showed increased cell cycle progression. We demonstrated that the addition of IGFBP2 to differentiating PSCs resulted in a higher number of functional progenitors, identifying the supporting role of arterial cells play to the emergence of blood progenitors via IGFBP2 paracrine signalling.

## Introduction

The haematopoietic system develops early during gestation through so called hematopoietic “waves” of progenitors, arising from different anatomical region and giving rise to various progenitor and stem cells (Medvinsky and Dzierzak, 1996; Palis *et al*., 1999; Böiers *et al*.,2013; Hoeffel *et al*., 2015; Patel *et al*., 2022). Although many types of hematopoietic progenitors of the various lineages can now be successfully produced from iPSCs, efficient production of HSCs remains still a standing challenge. The precise mechanisms leading to the development of functional HSCs is yet to be completely defined posing a limitation on how to recapitulate it in vitro.

During embryonic development, HSCs emerge from specialised endothelial cells lying on the ventral region of the dorsal aorta in the posterior region of the embryo (Jaffredo *et al*., 1998; Zovein *et al*., 2008; Bertrand *et al*., 2010; Boisset *et al*., 2010).

Only a subset of the endothelial cells, known as hemogenic endothelium, is capable of generating hematopoietic stem and progenitor cells via endothelial to hematopoietic transition (EHT) (Ottersbach, 2019). During the EHT endothelial cells slow their cell cycle (Batsivari *et al*., 2017; Canu *et al*., 2020), round up and eventually detach to enter the circulation (Eilken, Nishikawa and Schroeder, 2009; Kissa and Herbomel, 2010). These profound phenotypical changes are accompanied by transcriptional remodelling whereby the expression of endothelial genes is gradually downregulated and the transcription of the hematopoietic program is initiated (Swiers *et al*., 2013). With the use of single cell transcriptomics a population of hemogenic endothelium specifically committed to the development of HSCs was recently identified within the developing human dorsal aorta (Zeng *et al*., 2019).

To explore the molecular control on the development of the hematopoietic system, and to address the difference with the in vitro system, we compared our own single cell transcriptomics analysis of in vitro derived hemogenic endothelium and early progenitors (Fidanza *et al*., 2020) to that of the HSC-primed human hemogenic endothelium (Zeng *et al*.,2019). To assess the role of the genes that were expressed at a lower level within in vitro-derived cells compared to in vivo, we developed a novel DOX-inducible CRISPR gene activation system. We employed scRNAseq to track the presence of guide RNAs and monitored the phenotypic effects of gene activation of the induced cell populations. With this experimental pipeline we identified a novel role for IGFBP2 in the control of cell cycle progression within the EHT process.

## Results

### Comparison of in vitro hematopoiesis to in vivo AGM identifies 9 differentially expressed transcription factors

We, and others, have shown that in vitro differentiation of human iPSCs closely resemble intraembryonic hematopoiesis (Sturgeon *et al*., 2014; Ng *et al*., 2016; Fidanza *et al*., 2020; Calvanese *et al*., 2022). To understand the molecular basis underlying the challenges associated with the production of definitive fully mature HSCs in vitro from differentiating iPSCs we compared our scRNAseq dataset (Fidanza *et al*., 2020) to that of cells derived in vivo from the human aorta-gonad-mesonephros (AGM) region (Zeng *et al*., 2019), where definitive HSCs develop. We integrated the transcriptomic data of in vitro derived endothelial (IVD_Endo) and hematopoietic cells (IVD_HPC) with that of arterial endothelial cells (aEC), arterial hemogenic cells (aHEC) and venous endothelial cells (vEC) derived from human embryos collected between the Carnegie stage 12 and 14 (Figure 1A). To identify possible target transcription factor that could be manipulated in vitro to improve iPSCs differentiation, we determined which genes were expressed in the aHEC at higher level compared to IVD_Endo and IVD_HPC. This strategy identified 9 transcription factors *RUNX1T1, NR4A1, GATA2, SMAD7, ZNF124, SOX6, ZNF33A, NFAT5, TFDP2* (Figure 1B). The expression of these genes was consistently high in aHEC, with some also high in the aEC, but low in both endothelial and hematopoietic IVD cells, except for TFDP2 which was expressed in IVD_HPCs (Figure 1C).

**Figure 1.**
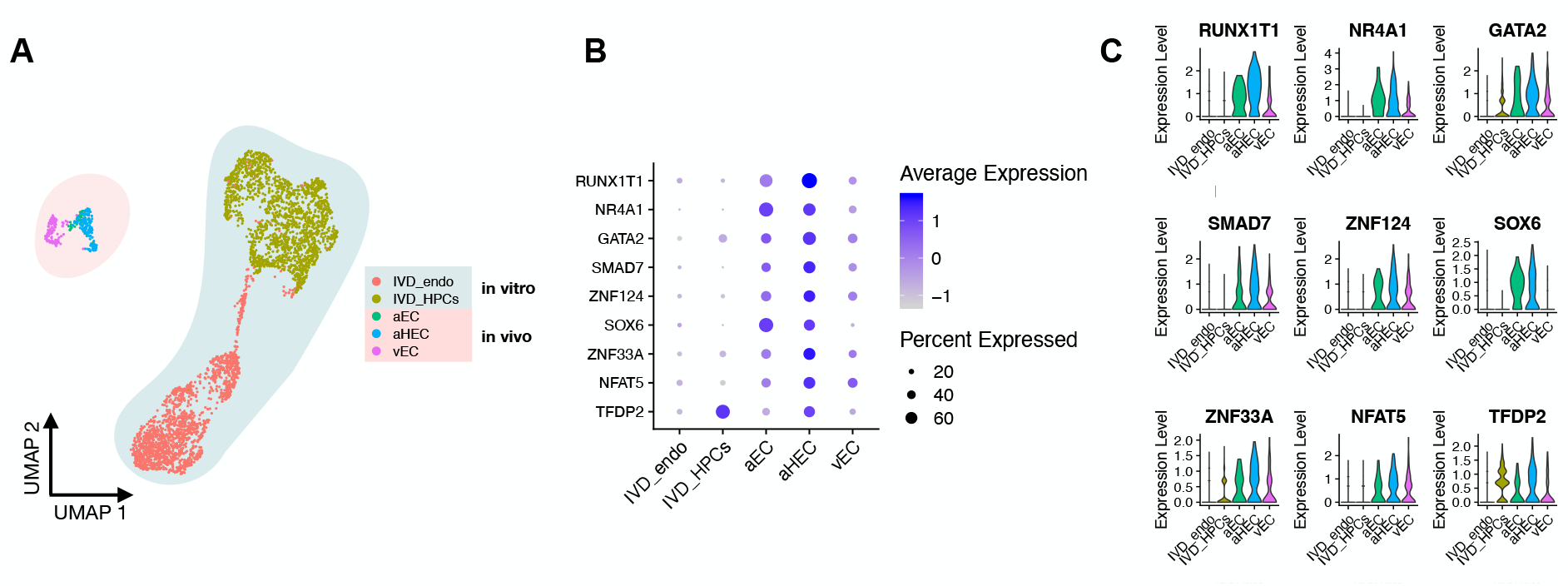
Comparison of in vitro hematopoiesis to in vivo AGM identifies 9 differentially expressed transcription factors. **A** - Integrative analysis of single cell transcriptome of in vitro derived hematopoietic (IVD_HPCs) and endothelial cells (IVD_Endo) and in vivo sample endothelial cells (venous, vEC; arterial, aEC; arterial hemogenic, HECs) from human embryos (CS12-CS14) visualised on UMAP dimensions. **B** - Target genes expression level showing higher expression in arterial hemogenic endothelium in vivo compared to in vitro derived cells. **C** - Violin plot visualising gene expression distribution of the target transcription factors visualised in the violin plot.

### Development of a gRNA-mediated DOX-inducible dCAS9-SAM activation system in human iPSCs

We previously developed an all-in-one SAM system that mediates the transcriptional activation of endogenous gene expression (Fidanza *et al*., 2017). To activate the nine target genes identified in this study we have since developed a novel DOX inducible SAM (iSAM) cassette targeted into the *AAVS1* locus of human iPSCs (Figure 2A). We first tested the iSAM plasmid by transient transfection of HeLa cells together with gRNAs directed to the *RUNX1C* transcriptional start site. We demonstrated that the activation of *RUNX1C* expression was correlated with the DOX concentration in a linear manner (Figure S1 A/B). To verify the activation in human PSCs we employed a RUNX1C-GFP human embryonic stem cell (hESC) reporter cell line and this strategy also allowed us to study gene activation at single cell resolution (Figure S1 C-G). As predicted, the level of expression of the mCherry tag within the iSAM cassette was proportional to the concentration of DOX (Figure S1 D-E) and to the number of cells in which the RUNX1C gene had been activated, as assessed by the presence of RUNX1C-GFP+ cells (Figure S1 D-F). Furthermore, the level of expression of RUNX1C expression, as measured by the mean fluorescence intensity (MFI) of the RUNX1C-GFP reporter, also correlated with the concentration of DOX (Figure S1 G). We then tested the iSAM cassette in our iPSC line (SFCi55) (Figure 2B-D). Only when iPSCs were transfected with iSAM and the gRNA for RUNX1 in presence of DOX, the expression of the *RUNX1C* gene was detected (Figure 2B). RUNX1 protein was also detected by immunocytochemistry in transiently transfected iPSCs (Figure 2C).

**Figure 2.**
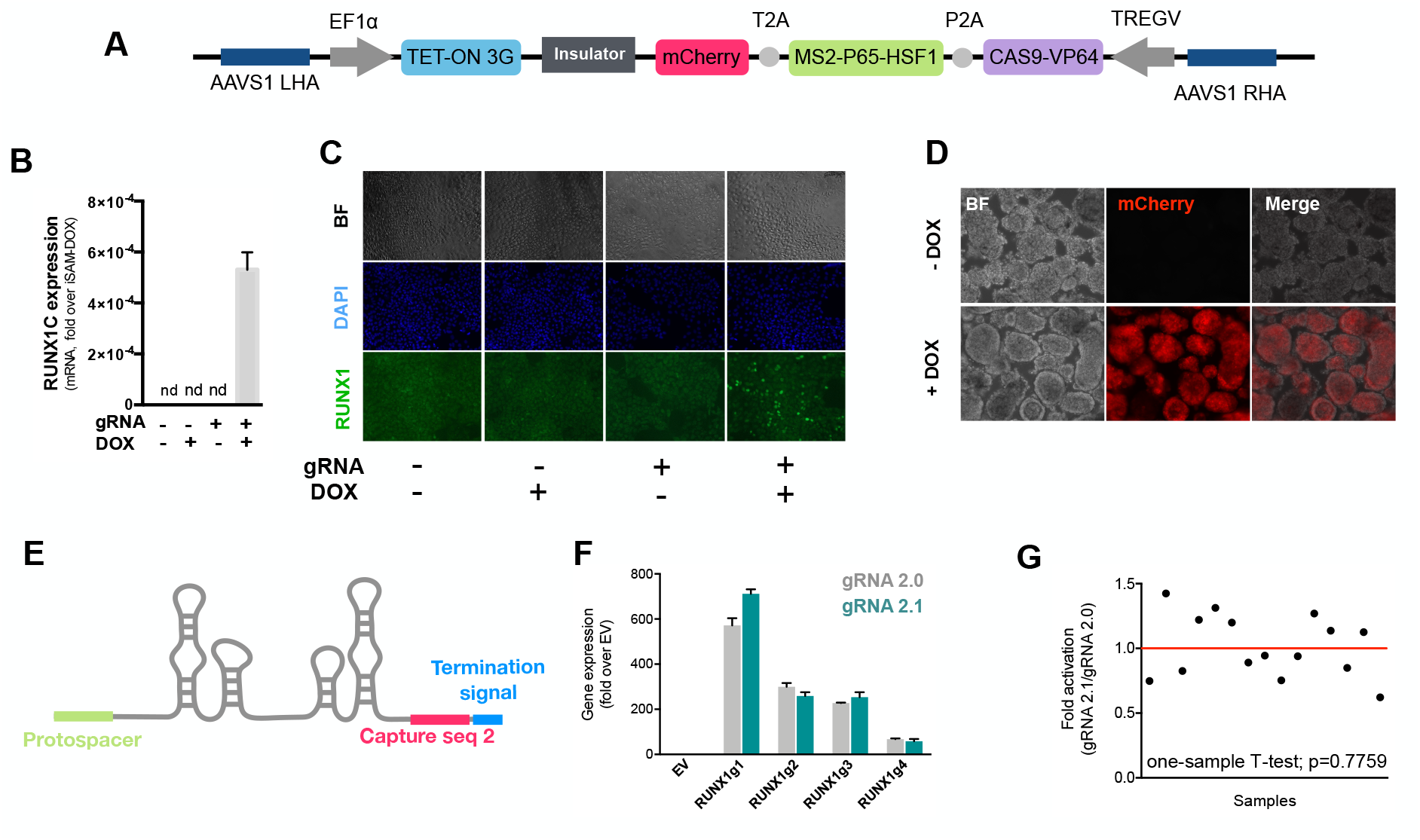
The inducible iSAM cassette successfully mediates activation of endogenous gene expression upon DOX induction. **A** - Schematic of the iSAM cassette containing the TET-on system under the control of EF1α and dCAS9-P2A-MS2-p65-HSF1-T2A-mCherry under the rTTA responsive elements, separated by genetic silencer and flanked by AAVS1 specific homology arms. **B** - *RUNX1C*gene expression activation after transient transfection of the iSAM plasmid and gRNAs in presence or absence of DOX in human iPSC line. **C** - RUNX1 protein expression upon iSAM activation after transient transfection of the iSAM plasmid and gRNAs with DOX in human iPSC line detected by immunostaining. **D** Expression of the iSAM cassette reported by mCherry tag during the differentiation protocol, the representative images (bright field – BF, and fluorescence) show embryoid bodies at day 3 of differentiation. **E** - Schematic of the gRNA 2.1 containing the capture sequence for detection during the scRNAseq pipeline. **F** - *RUNX1C* gene activation level obtained using either the gRNA 2.0 or 2.1 backbone. **G** - Statistical analysis of the gRNAs activation level showing no significant variation following addition of the capture sequence (triplicate for each of the 4 different gRNAs).

We then targeted the iSAM cassette into the *AVVS1* locus using a Zinc Finger Nuclease (ZFN) strategy (Yang *et al*., 2017; M. Lopez-Yrigoyen *et al*., 2018). iPSC clones that had specifically integrated the iSAM cassette into the *AAVS1* locus were validated by genomic PCR and sequencing. The AAVS1 locus has been reported to be a safe harbour site that is resistant to epigenetic silencing and indeed we had previously demonstrated that transgenes inserted into the *AVVS1* locus under the control of the constitutively active CAG promoter was efficiently expressed both in undifferentiated and in differentiated iPSCs (Yang *et al*.,2017; Martha Lopez-Yrigoyen *et al*., 2018; Lopez-Yrigoyen *et al*., 2019). However, we noted a dramatic reduction in the number of cells expressing the mCherry tag in undifferentiated iSAM iPSCs upon DOX induction after the iSAM line had been maintained for several passages, indicating transgene silencing of the rTTA DOX-inducible cassette (Supp Figure 1H). To overcome this problem, we treated the iSAM iPSC line with an inhibitor of histone deacetylases (HDACs), sodium butyrate (SB), known to have no adverse effect on iPSCs maintenance (Kang *et al*., 2014; Zhang, Xiang and Wu, 2014). A short 48 hours treatment resulted in a significant increase the number of mCherry+ cells upon DOX induction, proportional to the SB concentration (Figure S1I). We therefore maintained the iSAM iPSCs in the presence of SB and this fully restored the inducibility of the transgene with virtually all cells expressing mCherry in the presence of DOX (Figure S1J). Importantly, we noted no detrimental effect of SB on iPSC self-renewal nor on their haematopoietic differentiation capacity.

To test the effect of activating the 9 target genes on the transcriptomes of differentiating cells, we engineered the gRNAs so they could be detected within the single cell RNA sequencing pipeline (Replogle *et al*., 2020). To this end we inserted a capture sequence just before the termination signal to avoid any alteration in the secondary structure of the loops thus preserving the binding of the synergistic activators of the SAM system to the stem loops of the gRNA. Of the two capture sequences available we decided to use the one that was predicted to result in fewer secondary structure alterations and this new gRNA was named 2.1 (Figure 2E). We compared the activation level achieved with the new 2.1 gRNA to that of the original 2.0 backbone using various gRNAs targeting *RUNX1C* (Figure 1F). These results convincingly demonstrate that the addition of the capture sequence in the gRNA 2.1 does not alter the level of endogenous gene activation that could be achieved (Figure 1G).

### CRISPR activation results in expansion of arterial cells in association with higher hematopoietic progenitors’ potential

We designed 5-7 gRNAs to target the 200bp upstream of the transcriptional start sites of each of the 9 target genes identified by the comparison to the human AGM dataset. We subcloned these 49 gRNAs (Table 1) into the gRNA 2.1 backbone, and packaged them into lentiviral particles, (herein referred to as the AGM library) as well as a non-targeting (NT) gRNA. The iSAM iPSC line was infected to generate iSAM_AGM and iSAM_NT iPSCs. Cells were selected in puromycin, and their integration in the genome was confirmed by PCR and sequencing. The iSAM_AGM and iSAM_NT iPSCs were then differentiated in 3D embryoid bodies (EBs) until day 8, then dissociated and analysed by flow cytometry for the expression of endothelial and arterial markers, CD34 and DLL4, respectively. DOX was added from day 0 to both cell lines to be able to distinguish between the effect of DOX alone and that of target gene activation by the gRNAs (Figure 3A). Although DOX alone resulted in increase of CD34+DLL4+ cells (Figure S2A), the increase obtained with the AGM library was significantly higher (Figure S2A, Figure 3B), with a more than 3-fold expansion of phenotypical arterial cells. To verify that the increase in arterial endothelial cells was associated with a functional difference we isolated CD34+ cells using magnetic beads and cocultured 20000 cells on OP9 supportive stromal cells for 7 days, in the presence of the same differentiation cytokines. After one week, half of the OP9 cocultured cells were plated into colony forming assays and scored 14 days later. These results showed that the activation of the target genes in the iSAM_AGM in presence of DOX led to an increased number of CFU-E an CFU-GM and a reduction of CFU-M (Figure 3C), supporting the idea that activation of these genes in differentiating iPSCs resulted in a change in the types of haematopoietic progenitors produced.

**Figure 3.**
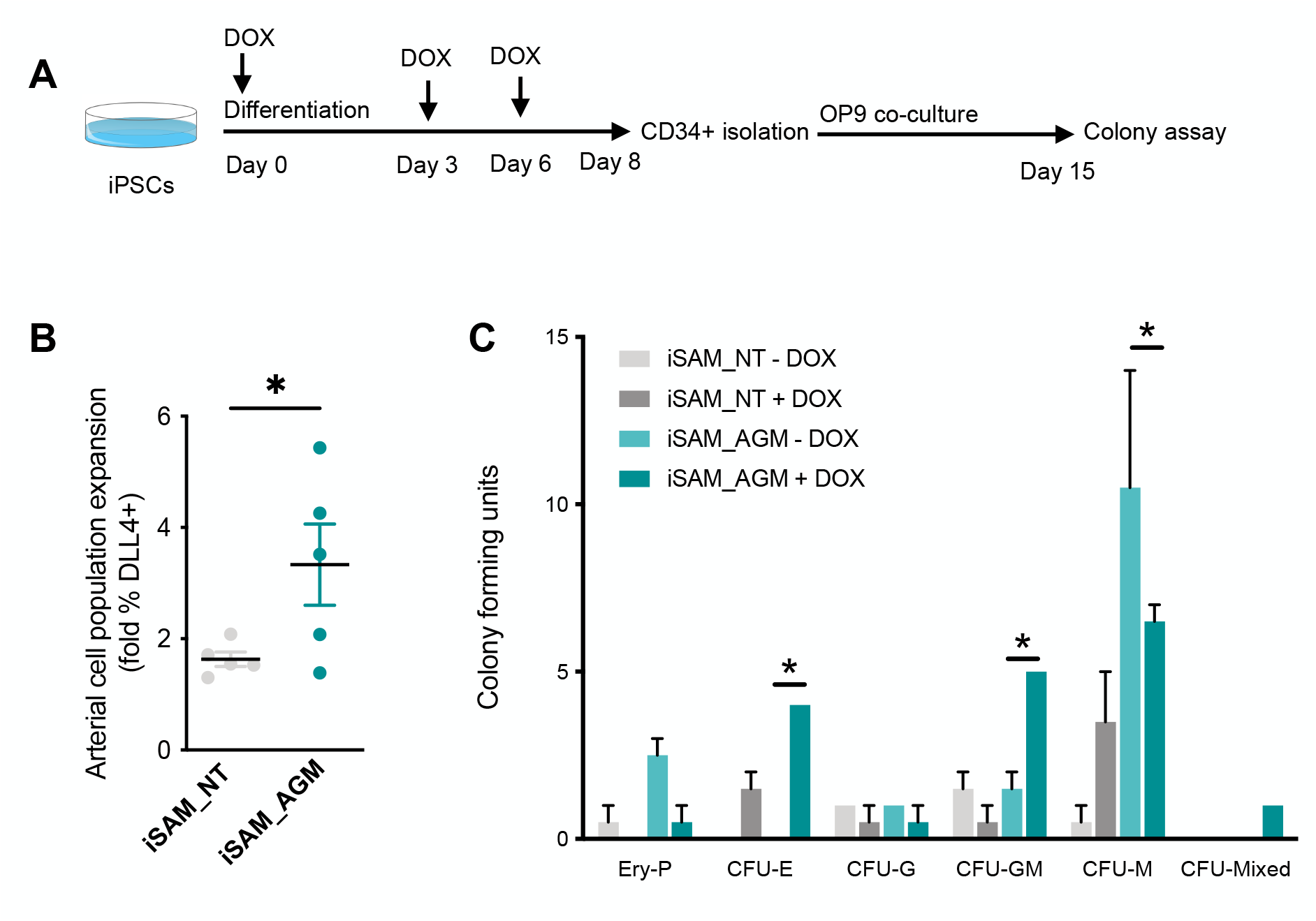
CRISPR activation results in expansion of arterial cells in association with higher clonogenic potential. **A** - Schematic of the differentiation protocol of the with activation of the target genes, used for both the control line iSAM_NT (containing the non-targeting control gRNA) and iSAM_AGM (containing the gRNAs for the target genes) **B** - Expansion of the arterial population assessed by the membrane marker expression of DLL4+ following targets’ activation, quantified by flow cytometry at day 8 of differentiation (Data are normalised on the iSAM_NT + DOX sample, * p = 0.0417 paired t-test). **C** - Colony forming potential of the suspension progenitor cells derived from the two lines treated with or without DOX following OP9 coculture activation, data show the colony obtained for 10^4^ CD34+ input equivalent (* p<0.05, Tukey’s two-way ANOVA).

### Single Cell RNA sequencing in combination with CRISPR activation identify arterial cell type expansion in association with activation of the 9 target genes and increased expression of *IGFBP2*

To analyse the transcriptional changes that were induced by the activation of the target genes, we differentiated the iSAM iPSCs and subjected them to single-cell RNA sequencing using the 10X pipeline. After 10 days of differentiation in the presence or absence of DOX, we FAC-sorted live CD34+ cells from iSAM_AGM and the iSAM_NT iPSCs (Figure 4A). Following data filtering of low-quality cells, we selected cells in which the gRNAs expression was detected (Figure 4B). To verify that our approach activated target genes we assessed the expression profile of these genes in the different libraries. All the target genes appeared to be expressed at a higher level in the iSAM_AGM compared to the iSAM_NT library (Figure 4C), as expected. Interestingly, many of these genes appeared downregulated upon DOX induction in the control iSAM-NT cell line but this effect was counteracted by the target gene activation in iSAM_AGM cells (Figure S2B). To study the effect of the genes’ activation on the cell types we performed clustering analysis and detected a total of 7 clusters (Figure 4B). High level of *DLL4* expression was detected in the arterial cell cluster while high levels of hemogenic-markers such as *RUNX1* and *CD44*, were detected in other clusters typed as hemogenic 1 and hemogenic 2 (Figure 4B, D). To understand the effect of the activation, we looked at the representation of the various clusters in the different libraries and we noticed a significant expansion of the arterial cluster in the DOX-induced iSAM-AGM library (Figure 4E). This is entirely in keeping with the expansion of DLL4+ cells that we had detected by flow cytometry (Figure 3B). We then compared the expression profile of these arterial cells between the different activation libraries, and we obtained a list of genes upregulated upon activation of the targets. One of these, *IGFBP2* was expressed at significantly higher levels in the iSAM_AGM library in presence of DOX compared to the others (Figure 4F), and this was associated with a significant enrichment of the RUNX1T1 specific gRNAs (23.67 average log2 fold change, 4.11 e^-07^ adjusted p-value).

**Figure 4.**
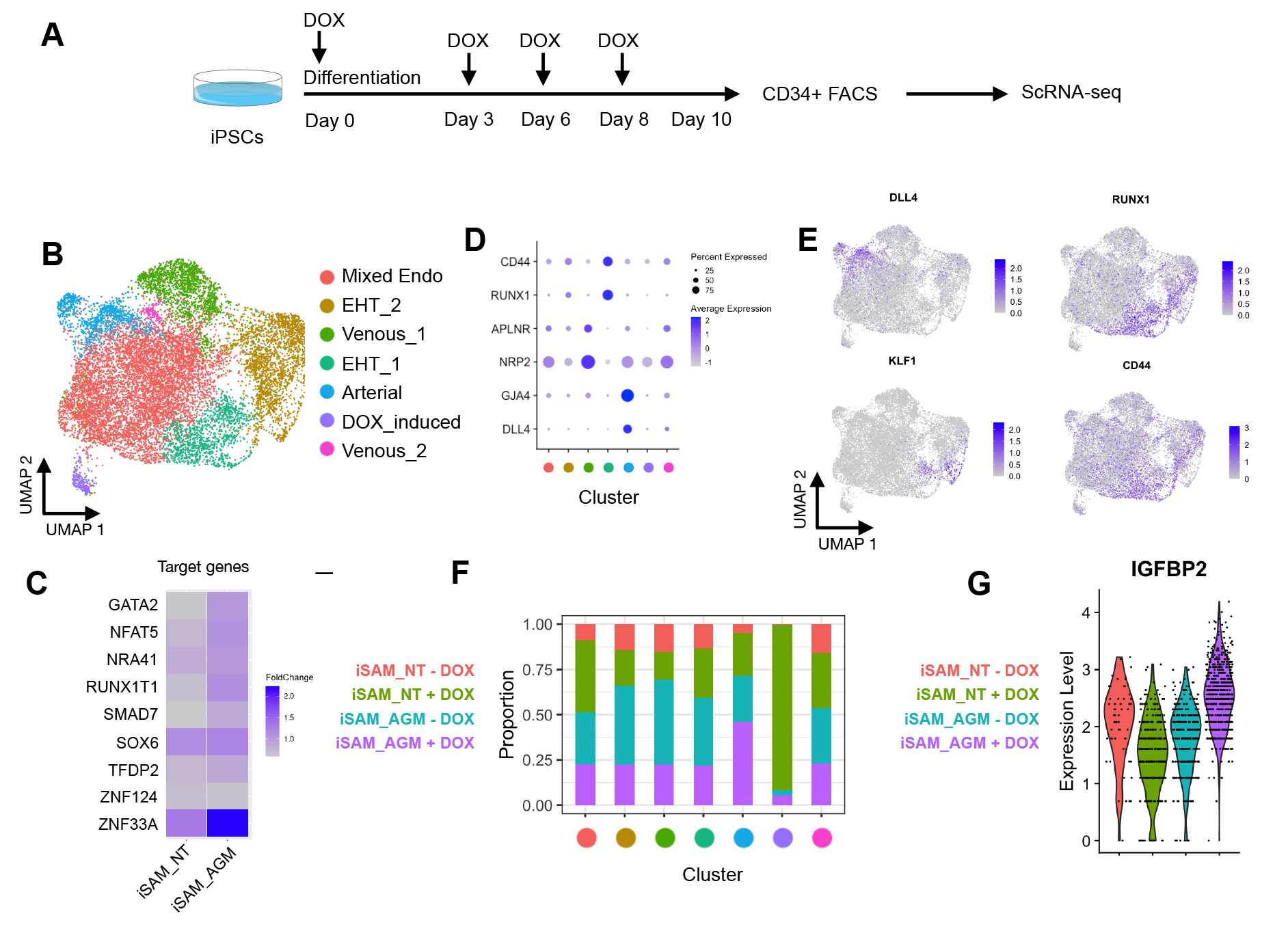
Single Cell RNA sequencing in combination with CRISPR activation of the 9 target genes identify arterial cell type expansion. **A** - Schematic of the single-cell RNAseq experiment in combination with the activation of the target genes. **B** - Dimension reduction and clustering analysis of the scRNAseq data following activation, filtered on cells where the gRNA expression was detected. **C** - Gene expression profile of target genes following target genes’ activation, heatmap shows the expression level of the target genes in the iSAM_NT and iSAM_AGM treated with DOX following normalisation on the -DOX control. **D** - Arterial (*GJA4, DLL4*), venous (*NRP2, APLNR*) and hemogenic marker (*CD44*, *RUNX1*) expression distribution in the clusters indicated by the colour. **E** - Expression distribution visualised on the UMAP plot showing the location of arterial cells marked by *DLL4*, and hemogenic endothelium marked by *CD44* and *RUNX1*, and hematopoietic priming marked by *KLF1*.**F** - Contribution of the different libraries to the clusters showing that arterial cell cluster is overrepresented in the iSAM_AGM treated with DOX, compared to the other libraries. **G** - Violin plot of *IGFBP2* expression profile in arterial cells obtained from the different conditions.

### IGFBP2 addition to the in vitro differentiation leads to a higher number of functional hematopoietic progenitor cells

IGF Binding Protein 2, a member of the family of IGF binding proteins, is able to bind both IGF1 and IGF2 as well as bind to other extracellular matrix proteins, for which it needs to be secreted by the cells. To test if the increased frequency of functional hematopoietic progenitors was due to paracrine signaling from the arterial cells, we supplemented the media with IGFBP2 at 100ng/ml from day 6, after the induction of endothelial cells differentiation (Figure 5A). To explore the role of IGFBP2 we employed the parental iPSCs line, SFCi55 from which the iSAM line was derived. We isolated CD34+ cells at day 8 and co-cultured them on OP9 cells in presence of IGFBP2 for an additional week, then tested them using colony forming unit (CFU) assays. The cells treated with IGFBP2 showed a significant increase in the total number of haematopoietic CFU colonies. (Figure 5B). To assess whether IGFBP2 also had a paracrine effect on the production of arterial cells themselves, we analyzed the cells within EBs at day 8, but no difference was detected in the number of DLL4+ cells in presence of IGFBP2 (Figure 5C). We then focused on the characterization of the cells derived from the CD34+ cells after coculture with the OP9. Our results showed a comparable distribution of CD34+, CD43+ and CD45+ cells population (Figure S2C-D), but a significant expansion of all these populations (Figure 5D). IGFBP2 has been previously reported to control HSCs cell cycle and support their survival ex-vivo by inducing proliferation (Huynh *et al*., 2011). Because cell cycle is tightly regulated during the EHT process both in vivo (Batsivari *et al*., 2017; Fadlullah *et al*., 2022) and in vitro (Canu *et al*., 2020), we explored the cell cycle stage of cells undergoing EHT in our dataset. EHT cells were subset from the initial dataset according to the expression of hemogenic markers such as RUNX1 and CD44. We performed pseudo-temporal ordering of the cells (Figure 5E) and looked at their cell cycle stage along pseudotime (Figure 5F). These analyses showed that cell undergoing EHT progress from G1 to renter the cell cycle in S and G2/M (Figure 5F). This observation was also in accordance with GO analysis of the clusters EHT_1 and EHT_2, showing an enrichment for ribosome associated GO in the EHT_1 and cell cycle and DNA replication associated GO in EHT_2, reflecting the progression of cells along the cell cycle stages (Figure S3A-B). In addition, the iSAM_AGM cells showed a higher number of cycling cells in S and G2M with a consequent reduction of G1 cells (Figure 5G-H). To uncouple the effect of IGFBP2 on cell cycle during the EHT from that on the progenitor population, we analysed the cell cycle profile of suspension hematopoietic cells obtained from OP9 cocultured in the presence of IGFBP2. This showed no differences on the cell cycle distribution of hematopoietic progenitors (Figure S3C) indicating that the expansion of hematopoietic progenitors is not a consequence of their increased cycling but rather the effect of IGBP2 during their emergence.

**Figure 5.**
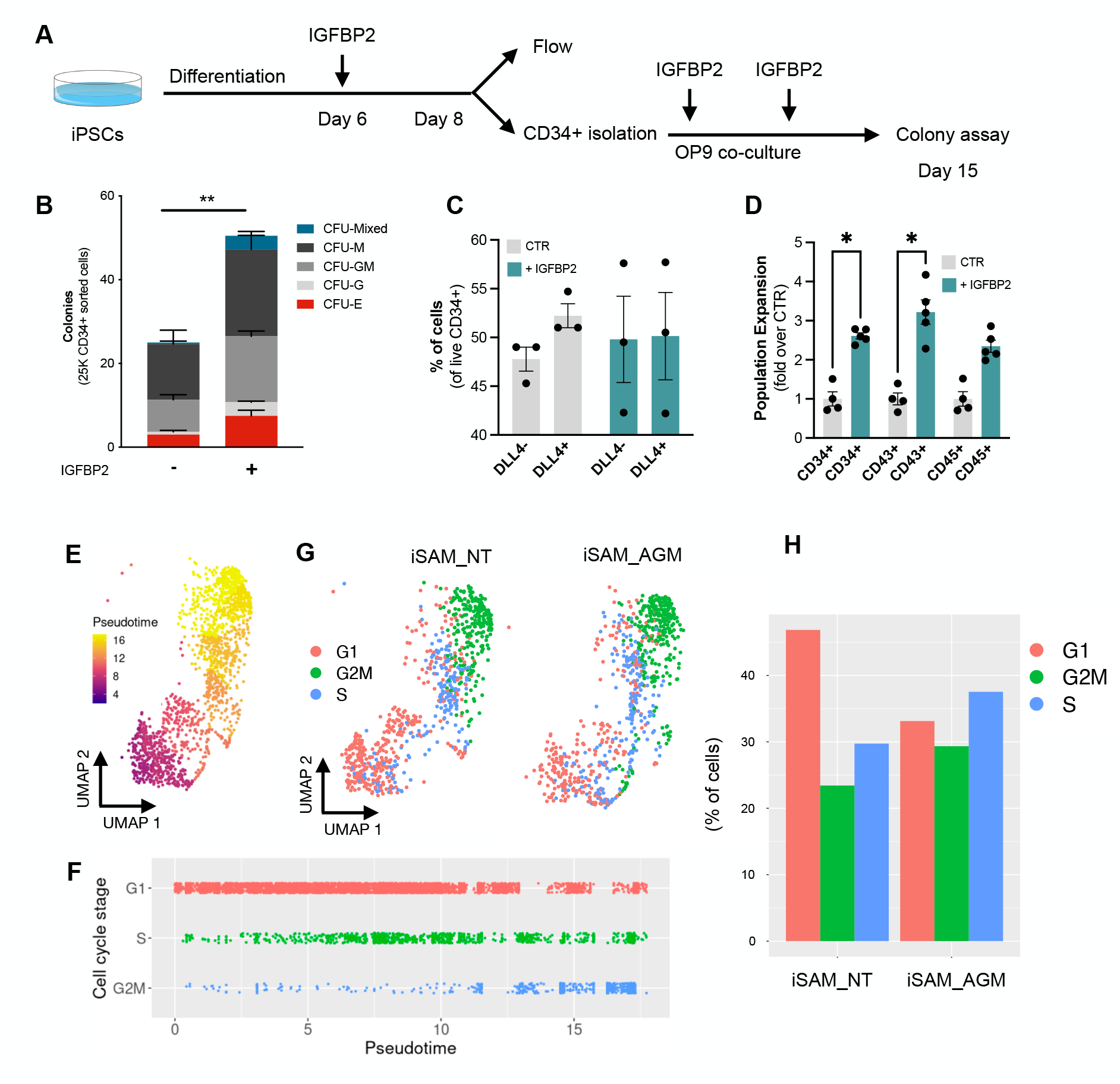
IGFBP2 addition to the in vitro differentiation leads to a higher number of functional hematopoietic progenitor cells. **A** - Schematic of the IGFBP2 functional validation experiment. **B** - Number of hematopoietic colonies obtained after coculture on OP9 in presence or absence of IGFBP2 (** p=0.0080, Sidak’s Two way ANOVA). **C** - Percentage of arterial cells differentiation analysed by flow cytometry for DLL4 in day 8 EBs. **D** - Expansion of hematopoietic progenitors analysed using markers’ expression on suspension progenitors derived after coculture of CD34+ cells onto OP9 support (data are expressed as fold over the CTR in the absence of IGFBP2 (* p<0.02, Sidak’s Two way ANOVA). **E** - Pseudotemporal ordering in the EHT cells’ subset showing the progression of cells from EHT_1 to EHT_2 cluster. **F** - Cell cycle stage ordered along the pseudotime axis during the process of the EHT. **G** - Cell cycle stages in the two libraries iSAM_NT and iSAM_AGM treated with DOX projected on the UMAP plot. **H** - Quantification of the different cell cycle stages in the iSAM_NT and iSAM_AGM cells treated with DOX.

Taken together these data suggests that the increased number of functional hematopoietic progenitors detected upon activation of the target genes could be explained by the enhanced production of IGFBP2 in arterial cells that support the EHT process by promoting re-entry in the cell cycle.

## Discussion

The complexity and dynamism of the developmental hematopoiesis in vivo has imposed challenges in accurately reproducing the process in vitro. Here we identified differences in the expression of nine transcription factors between the two systems and developed a novel CRISPR-activation system to explore the downstream consequences of their activation on the emergence of definitive hematopoietic progenitor cells. Using this approach, we identified the supportive role of non-hemogenic endothelial cells via paracrine signaling of IGFBP2 during the endothelial to hematopoietic transition.

When we compared endothelial cells derived in vitro from hiPSCs to those primed to give rise to HSCs, at the time point when the early commitment takes place in the human embryo, we identified 9 transcription factors that were expressed at lower level in vitro. While some of these have been associated with blood cell development such as GATA2 (Ling *et al*., 2004; de Pater *et al*., 2013; Castaño *et al*., 2019), SMAD7 (McGarvey *et al*., 2017), NR4A1(McGarvey *et al*., 2017), and SOX6 (McGrath *et al*., 2011), the others, RUNX1T1, ZNF124, ZNF33A, NFAT5 and TFDP2 have not been previously associated with haematopoiesis. The addition of capture sequence to the gRNA backbone enabled their detection coincidentally with the single-cell transcriptome and this allowed us to demonstrate that that RUNX1T1 gRNAs were significantly enriched in cells within the expanded arterial cluster. RUNX1T1, also known as ETO, has been associated with the t(8;21) chromosomal translocation that results in the generation of the leukemic fusion protein, AML1/ETO (Rejeski, Duque-Afonso and Lübbert, 2021). *RUNX1T1* expression has been recently detected in transcriptomic analysis of the human AGM region (Zeng *et al*., 2019; Calvanese *et al*., 2022), but its precise role during the ontogeny of the blood system has not been elucidated.

Here we show that enrichment of the RUNX1T1 gRNAs was associated with high expression of *IGFBP2*, encoding a member of the IGF binding protein family. *IGFBP2* KO mice show increased expression of cell-cycle inhibitors and HSC apoptosis, implicating IGFBP2 as a modulator of HSCs cell cycle and survival (Huynh *et al*., 2011). More recently *IGFBP2* was reported as being highly expressed in the human AGM region at CS14 when HSCs are emerging (Calvanese *et al*., 2022), supporting a possible role during developmental hematopoiesis. In this study we show that the addition of IGFBP2 recombinant protein in our in vitro model of EHT results in the emergence of an increased number of functional hematopoietic progenitors. This higher number was not associated with increased cell cycle division in the progenitors per se but rather to the faster progression towards G2/S/M in the hemogenic endothelial cells undergoing EHT. Cell cycle control has been identified to be an important mediator of EHT; first the cell cycle slows down to allow the intense cell remodeling that leads to the acquisition of the hematopoietic potential to then restart ensuring the proliferation of blood progenitors (Batsivari *et al*., 2017; Canu *et al*.,2020; Fadlullah *et al*., 2022). The molecular control of this process is still unclear and here we demonstrate that role of IGFBP2 as mediator of the process and propose that RUNX1T1 might be regulating its expression. Because RUNX1T1 lacks a DNA binding domain, its direct involvement in the regulation of IGFBP2 expression must require the involvement of other cofactors that are yet to be identified.

The supportive role of the endothelial niche in the development of HSCs has been studied in vivo (Hadland *et al*., 2015, 2022; Crosse *et al*., 2020) and exploited in vitro to support HSCs emergence (Sandler *et al*., 2014; Hadland *et al*., 2022). Although many signalling molecules have been associated with HSCs development, this is the first indication that IGFBP2 could be one of the supportive mediators of the EHT by modulating the cell cycle.

This study demonstrates that the combination of CRISPR mediated activation of target genes with single cell transcriptomic analysis in differentiating PSCs can be a powerful approach to model human embryonic development. The fine epigenetic manipulation of the transcription permits the study of target gene sets simply by adding specific gRNAs and the strategy can be readily applied to any cell lineage of interest. Our findings underline the importance of mimicking the functional heterogeneity of developing tissues in vitro to recapitulate with accuracy the complex processes that occur in the developing embryo.

## Methods

### Pluripotent Stem Cells maintenance

hPSCs were maintained in vitro in StemPro hESC SFM (Gibco) with bFGF (R&D) at 20 ng/ml. Wells were coated with Vitronectin (ThermoFisher Scientific) at least 1 hour before plating and cells were passaged using the StemPro EZPassage tool (ThermoFisher Scientific). Media change was performed every day and cells passaged every 3-4 days at a ratio of 1:4.

### Transfection

iPSCs SFCi55 and hESCs RUNX1-GFP were plated at 3 × 10^5^ cells per a well of a 6 well plate and reverse transfected with 2 μg of DNA using the Xfect Transfection reagent (Clontech) and analyzed 2 days later.

HeLa cells were cultured in Dulbecco’s Modified Eagle Medium/Nutrient Mixture F-12 (DMEM/F12) with Glutmax and 5% FCS (Gibco) and passaged every few days, at a ratio of 1:6. HEL were cultured in Iscove’s Modified Dulbecco’s Medium (IMDM) with 10% FCS (Gibco) and passaged every few days, at a ratio of 1:4. 2 × 10^5^ cells were plated, transfected at 6–8 hours with 0.75 μg of DNA using Xfect Transfection reagent (Clontech) and then analysed 2 days after.

### Immunocytochemistry

Cells were fixed in 4% PFA in PBS at room temperature for 10’, permeabilized in PBS-T (0.4% Triton-X100) for 20’ and blocked in PBS-T with 1% BSA and 3% goat serum for 1 hour. Primary antibodies were incubated in blocking solution over night at 4 °C (RUNX1 1:200 - ab92336, Abcam). Cells were then washed in PBS-T and incubated with secondary antibodies for 1 hour at room temperature (donkey α-rabbit 1:200 - A-11008 - Thermo Scientific). Cells were washed in PBS-T and counterstained with DAPI. Images were generated using the Zeiss Observer microscope.

### Gene expression analysis

Total RNA was purified using the RNAeasy Mini Kit (Qiagen) and cDNA synthesized from 500 ng of total RNA using the High Capacity cDNA synthesis Kit (Applied Biosystem). 2 ng of cDNA were amplified per reaction and each reaction was performed in triplicate using the LightCycler 384 (Roche) with SYBR Green Master Mix II (Roche). A melting curve was performed and analyzed for each gene to ensure the specificity of the amplification. *β-Actin* was used as reference genes to normalize the data (Fidanza *et al*., 2017).

### Pluripotent Stem Cells differentiation to hematopoietic progenitors

hPSCs were differentiated in a xeno-free composition of SFD medium (Fidanza *et al*., 2020), BSA was substituted with human serum albumin, HSA (Irvine-Scientific). Day 0 differentiation medium, containing 10 ng/ml BMP4 was added to the colonies prior cutting. Cut colonies were transferred to a Cell Repellent 6 wells Plates (Greniner) to form embryoid bodies and cultured for two days. At day 2 media was changed and supplemented with 3 μM CHIR (StemMacs). At day 3, EBs were transferred into fresh media supplemented with 5 ng/ml bFGF and 15 ng/ml VEGF. At day 6 media was changed for final haematopoietic induction in SFD medium supplemented with 5 ng/ml bFGF, 15 ng/ml VEGF, 30 ng/ml IL3, 10 ng/ml IL6, 5 ng/ml IL11, 50 ng/ml SCF, 2 U/ml EPO, 30 ng/ml TPO, 10 ng/ml FLT3L and 25 ng/ml IGF1. From day 6 onward, cytokines were replaced every two days.

### CD34 isolation

CD34+ cells were isolated using CD34 Magnetic Microbeads from Miltenyi Biotec, according to their manufacturing protocol. Briefly, Embryoid bodies were dissociated using Accutase (Life Techonologies) at 37°C for 30’. Cells were centrifuged and resuspended in 150 μl of PBS + 0.5% BSA + 2mM EDTA with 50 μl Fcr blocker and 50 μl of magnetic anti-CD34 at 4°C for 30’. Cells were washed using the same buffer and transferred to pre-equilibrated columns, washed three times and eluted. After centrifugation, cells were resuspend in SFD media, counted and plated for OP9 coculture.

### OP9 coculture and colony assay

OP9 cells were maintained in a-MEM supplemented with 20% serum (Gibco) and sodium bicarbonate (Gibco) and passaged with Trypsin every 3-4 days. The day before the co-colture, 45.000 OP9 cells were plated for each 12 well plates’ well in SFD media. The day of the co-culture the 20.000 iSAM cells or 25.000 SFCi55 or H9 were plated in each well and culture in SFD media supplemented with 5 ng/ml bFGF, 15 ng/ml VEGF, 30 ng/ml IL3, 10 ng/ml IL6, 5 ng/ml IL11, 50 ng/ml SCF, 2 U/ml EPO, 30 ng/ml TPO, 10 ng/ml FLT3L and 25 ng/ml IGF1 and 100ng/ml IGFBP2. Cytokines were replaced twice during one week of coculture. At the end of the coculture, cells were collected by Trypsin and half of the well equivalent was plated in 2 ml of methylcellulose medium (Human enriched H4435, Stemcell Technologies). Cells were incubated in the assay for 14 days and then scored.

### Methylcellulose assay

OP9 co-cultured hiPSCs progenitor cells were collected by Trypsin treatment and resuspend in SFD medium. Half well equivalent was seeded into 2 ml of methylcellulose medium (Human enriched H4435, Stemcell Technologies). Cells were incubated in the assay for 14 days and then scored.

### Flow cytometry staining and cell sorting

Embryoid bodies were dissociated using Accutase (Life Techonologies) at 37°C for 30’. Cells were centrifuged and resuspended in PBS + 0.5% BSA + 2mM EDTA, counted and stained at 10^5^ cells for a single tube. Cells were stained with antibodies for 30’ at room temperature gently shaking. Flow cytometry data were collected using DIVA software (BD). For the sorting experiments, cells were stained at 10^7^cells/ml in presence of the specific antibodies. Sorting was performed using FACSAria Fusion (BD) and cells were collected in PBS + 1% BSA. Data were analysed using FlowJo version 10.4.2.

### Flow cytometry antibodies

For flow cytometry 10^5^ cells per test were stained in 50 μl of staining solution with the following antibodies: CD34 Percp-Efluor710 (4H11 eBioscience, 1:100), CD34 Pe (4H11 eBioscience, 1:200), CD43 APC (eBio84-3C1, 1:100), CD45 FITC (2D1 ebioscience, 1:100), DLL4 Pe (MHD4-46 Biolegend, 1:200)

### iSAM plasmid generation

The iSAM plasmid was obtained by Gibson assembly of four fragments. The first fragment, the backbone, was a DOX-inducible AAVS1 targeted plasmid expresing an E6-E7-IRES-ZsGreen which was excised by BstBI and NdeI. The second fragment, one of the adapters, was derived from the UniSAM plasmid that we previously generated (Addgene #99866) by PCR with the following primer sets FW_aggggacccggttcgagaaggggctcttcatcactagggccgctagctctagagagcgtcgaatt, RV_ttcgggtcccaattgccgtcgtgctggcggctcttcccacctttctcttcttcttggggctcatggtggcc. The UniSAM cassette was obtained also from the UniSAM plasmid via digestion with BsrGI and BsiWI. Finally the last fragment consisting of another adapter for the Gibson was custom synthetised and contained overlapping sequences flanking a chicken b-globin insulator that we inserted to prevent cross-activation of the EF1α-promoter and the TRE-GV promoter driving the iSAM. Correct assembly was verified by Sanger sequencing. The plasmid will be deposited to Addgene (Pending submission), we will add the code upon receipt from Addgene.

### iSAM cell lines derivation

The iSAM plasmid was used together with ZNFs specific for the AAVS1 locus to mediate specific integration in SFCi55 human iPSCs line (Yang *et al*., 2017; Fidanza *et al*., 2020). Briefly, 10 μg of AAVS1-iSAM with 2.5 μg of each ZNFs, left and right, using Xfect (Takara) according to the manufacturer protocol. Cells were selected using Neomycin. Single clones were picked, amplified, and initially screened by mCherry expression upon DOX addition. Clones that expressed the fluorescent tag were screened for specific integration using PCR followed by Sanger sequencing for the correctly integrated clones. 100 ng of genomic DNA was amplifies using theEmeraldAmp^®^ MAX HS Takara and specific primers sets (Table1).

### Capture sequencing addition to the gRNA backbone

The Capture sequence 2 was added to the gRNA_Purp_Backbone (Addgene #73797) by PCR. Briefly, the capture sequence was added before the termination signal of the gRNA followed by a BamHI site using the following PCR primers: gRNA_FW gagggcctatttcccatgattcct, gRNA_Cap_RV aaaaaaggatccaaaaaaCCTTAGCCGCTAATAGGTGAGCgcaccgactcggtgcc. The gRNA backobone was replaced from the original plasmid via NdeI and BamHI digestion, followed by ligation of the PCR produced following the same digestion. Correct integration of the insert was verified by Sanger sequencing.

### AGM library preparation

sgRNA design was performed by selecting the top candidates for on-target and off-target score. Between 5 and 7 guides per gene were designed for *RUNX1T1, NR4A1, GATA2, SMAD7, ZNF124, SOX6, ZNF33A, NFAT5, TFDP2* using the CRISPRpick tool from the Broad Institute (https://portals.broadinstitute.org/gppx/crispick/public). All the guide variants were Golden Gate cloned with the gRNA 2.1 backbone according to the established protocol (Konermann *et al*., 2015). The 49 plasmids were pooled together in an equimolar ratio and the library prep was subsequently used to produce lentiviral particles with a second-generation production system. Briefly, the psPAX2 packaging plasmid, pMD2.G envelope, and the AGM vector library were co-transfected using polyethyleneimine (PEI) (Polysciences, Warrington, PA, USA) as previously detailed (Petazzi *et al*., 2020), Lentiviral particles-containing supernatants were harvested 48–72 h post-transfection, concentrated by ultracentrifugation and titered in hiPSCs cells.

### iSAM_AGM and iSAM_NT cell line derivation

The selected iSAM clone (3.13 internal coding) was infected with viral particles containing either the AGM library or the non targeting gRNA (NT) at a MOI of 10. The iSAM cells were plated the afternoon before at 7×10^6^ cells into a T125 in presence of 10 μM Rock Inhibitor (Merk) which was maintained until the day following the infection. Cells were infected in presence of 8μg/ml of Polybrene (Merk). Puromycin selection was initiated 36 hours post-infection and maintained during their culture until the beginning of the differentiation.

### Single Cell RNA sequencing

Embryoid bodies obtained from day 10 of differentiation were dissociated using Accutase (Life Techonologies) at 37°C for 30’. Cells were centrifuged and resuspended in CD34-Pe staining solution at a density of 10^7^/ml. CD34+/live/single cells were FAC-sorted in PBS + 0.1 % BSA. Cell viability was also confirmed by Trypan blue stain for accurate count. Around 15000 cells per sample were loaded into the 10X Chromium Controller and single cell libraries were obtained using the Chromium single cell 3’ Reagent Kits v3 (10XGenomics) according to manufacturer protocol. The four libraries were indexed using SI PCR primers with different i7 indexes to allow for demultiplexing of the sequencing data. RNA concentration was obtained using Quibit RNA HS (Thermo-Fisher). Quality of the obtained libraries were verified using LabChip GX (PerkinElmer). Libraries were sequenced using NextSeq 2000 technology (Illumina) at 50.000 reads/cell. Data were aligned to GRCh38 using the Cell Ranger dedicated pipeline (10XGenomics). Data filtering, dimension reduction, clustering analysis, differentially expressed genes and cell cycle analysis were obtained using Seurat R package (version 4.1.0)(Hao *et al*., 2021). Pseudotemporal ordering was performed using Monocle 3 R package (Cao *et al*., 2019). KEGG pathways was performed using ShinyGo(Ge, Jung and Yao, 2020). The code will be made available on Github prior to publication, the raw data will be submitted to Array Express and the browsable process data will be added to our website containing previous sequencing data at https://lab.antonellafidanza.com.

## Data availability

R code is available at https://github.com/afidanza/CRISPRa. Data will be deposited to ArrayExpress for the final peer-reviewed version. Plasmids will be deposited to Addgene before publication following peer-review.

## Author contribution

AF designed the study, performed experiments and bioinformatic analysis, wrote the paper and led the research. PP, TV, FPL performed experiments. AM and HT provided support to the experiments. NR performed bioinformatic analysis. LF and PM helped the design of the study and the research. All authors provided essential feedback on the experiments and to the manuscript.

## Acknowledgment

AF and LF acknowledge financial support from the Biotechnology and Biological Sciences Research Council; Grant S002219/1. AF was supported by a European Hematology Association Advanced Research Grant (EHA RAG 2021). TV and AM were supported from PhD studentships from the Medical Research Council (Precision Medicine) and College of Medicine and Veterinary Medicine, respectively. FPL was supported by a Erasmus+ Traineeship Program 2016/2017. PM acknowledges financial support from a PERIS program from the Catalan Government and a Retos collaboration project from the MINECO (RTC-2018-4603-1)

## Supplementary Figures

**Supplementary Figure 1.**
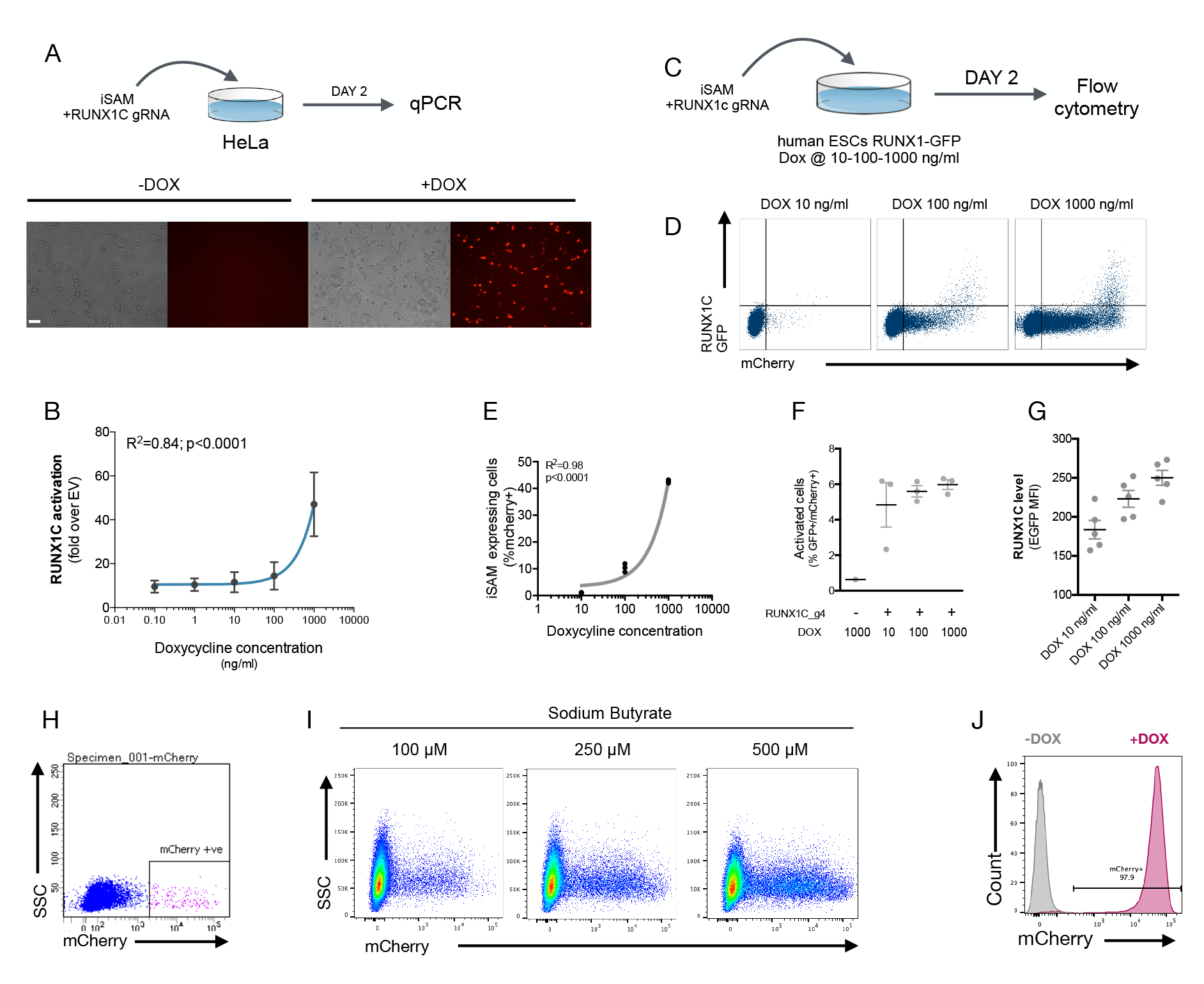
**A** – Schematic of the iSAM mediated activation of RUNX1C by transient transfection in HeLa cells with the iSAM vector and the RUNX1C gRNA; fluorescent microscopy demonstrating the expression of the mCherry tag. **B** – Linear regression of *RUNX1C* RNA expression in relation to the concentration of DOX added to HeLa cells. **C** - Schematic of the iSAM mediated activation of the hESCs RUNX1C-GFP reporter cell line by transient transfection of the iSAM vector and RUNX1C gRNA. **D** – Flow cytometry analysis of RUNX1C-GFP expression and mCherry in the hESCs RUNX1C-GFP reporter line, after exposure to different DOX concentration. **E** - Linear regression of the mCherry tag expression in relation to the concentration of DOX added to the hESCs RUN1C-GFP reporter line. **F** – Percentage of activated cells (GFP^+^mCherry^+^) in presence of different concentration of DOX. **G** – RUNX1C single cell expression level analysed by flow cytometry in hESCs exposed to different DOX concentration. **H** – Flow cytometry analysis of mCherry+ cells upon DOX addition in hiPSCs with iSAM targeted into the AAVS1 locus following maintenance. **I** – Flow cytometry analysis of the expression of the mCherry tag, in hiPSCs with iSAM targeted into the AAVS1 locus, following 48h Sodium Butyrate and DOX treatment at different concentration. **J** - Flow cytometry analysis of the expression of the mCherry tag upon DOX addition, in hiPSCs with iSAM targeted into the AAVS1 locus, maintained in presence of 500 μM Sodium Butyrate.

**Supplementary Figure 2.**
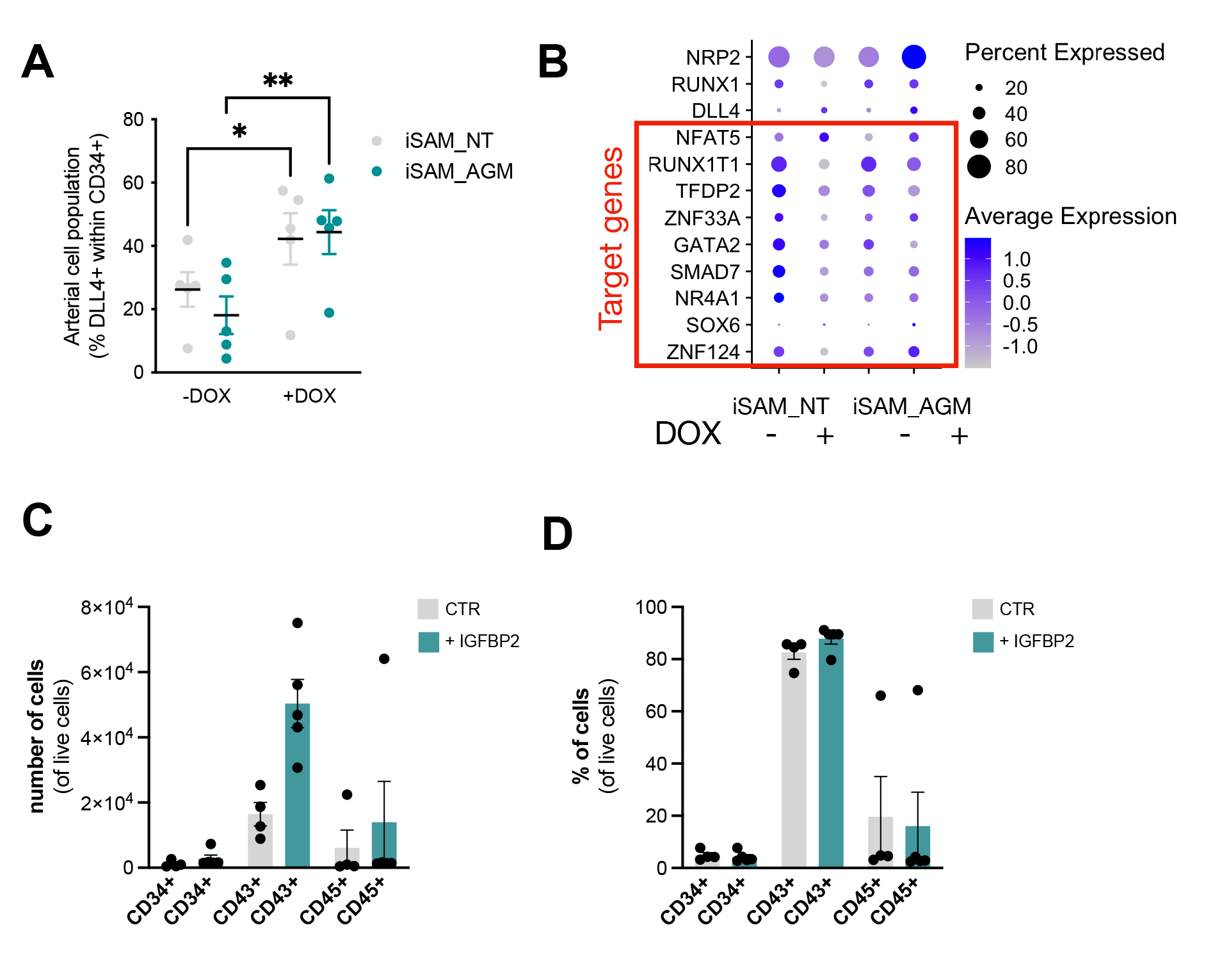
**A** – Expansion of the arterial population marker by membrane expression of DLL4+ following targets’ activation, quantified by flow cytometry at day 8 of differentiation (* p = 0.0190, ** p = 0.0011, Sidak’s Two-way ANOVA). **B** – Gene expression level of the target genes and additional markers (venous *NRP2*, hemogenic *RUNX1*, arterial *DLL4*) in the iSAM_NT and iSAM_AGM cell lines upon DOX addition. **C** - Number of hematopoietic cells expressing progenitors’ marker after OP9 coculture analysed by flow cytometry. **D** – Percentage of hematopoietic cells expressing progenitors’ marker after OP9 coculture analysed by flow cytometry out of total live cells.

**Supplementary Figure 3.**
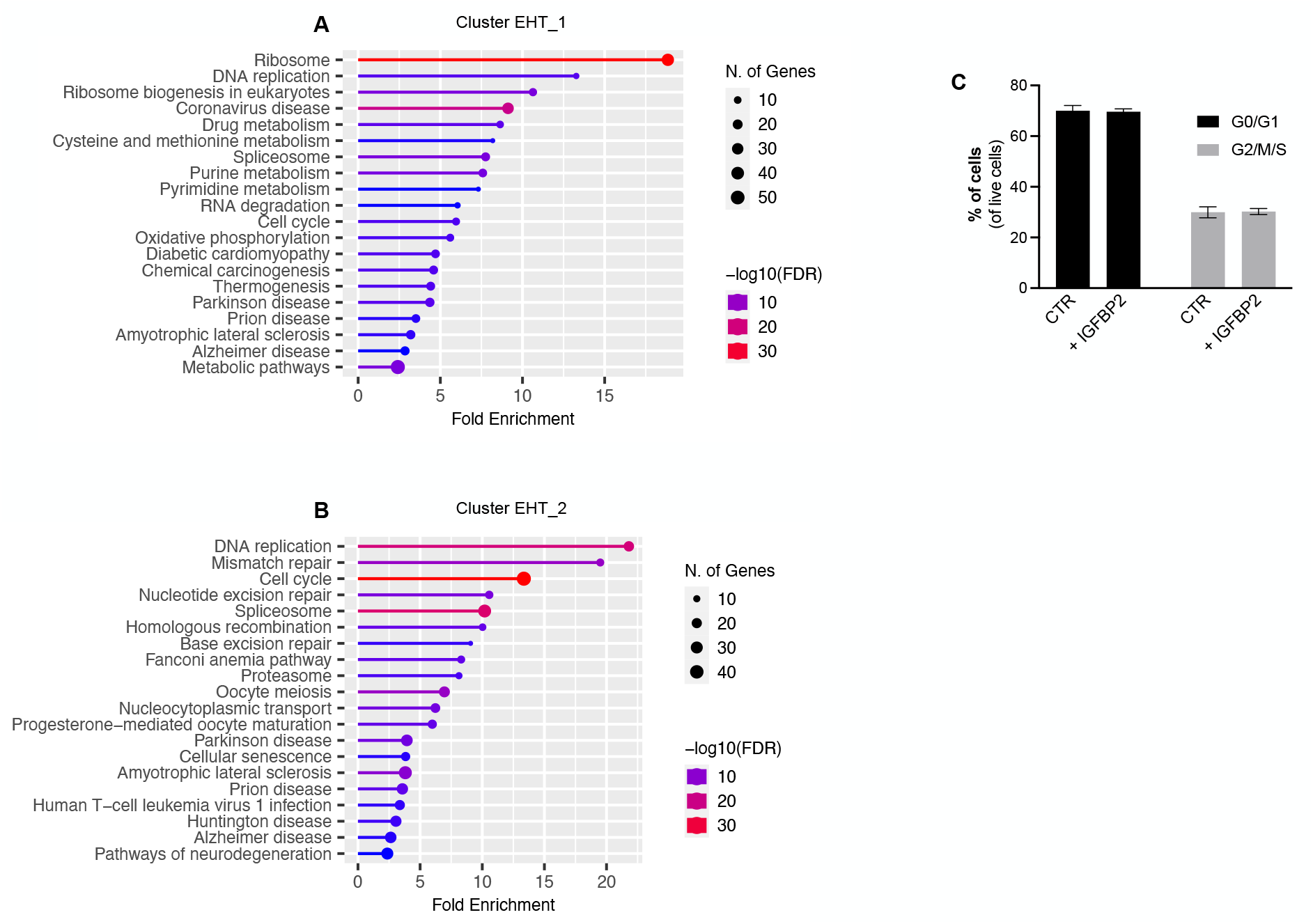
**A** – Gene Ontology analysis of marker genes of the cluster EHT_1. **B** – Gene Ontology analysis of marker genes of the cluster EHT_1. **C** – Flow cytometry analysis of the cell cycle analysis of suspension progenitor cells obtained following OP9 coculture of cells treated with IGFBP2 and control.

## Notes

### Competing Interest Statement

The authors have declared no competing interest.

